# Chromosomal-level genome assembly of *Populus adenopoda*

**DOI:** 10.1101/2023.07.11.548479

**Authors:** Shuyu Liu, Zeng Wang, Tingting Shi, Xuming Dan, Yulin Zhang, Jianquan Liu, Jing Wang

## Abstract

High-quality reference genomes for several species have promoted breeding and functional studies of poplar trees. By resequencing numerous accessions of these and closely related species, single nucleotide polymorphisms (SNPs) and small insertion/deletions (InDels) have been identified to assist in clarifying local adaptation and phenotypic diversification. A chromosome-level genome assembly for *P. adenopoda* was assembled based on Illumina and PacBio sequencing platforms, facilitated by Hi-C technology. The assembled genome size was about 383 Mb, with 99.70% of the contigs anchored to 19 pseudo-chromosomes, and a total of 33,505 protein-coding genes were annotated. This high-quality genome provided the genomic basis for the subsequent detection of various variants.

## Introduction

Trees comprise the main elements of the terrestrial landscape and are the basis for stability of ecosystems. Understanding the genetic diversity landscape of forest tree species from various spatial and temporal scale not only can inform current utilization, but also are important for the evaluation and conservation of wild germplasm resources. As one of the widespread and keystone forest tree species in China, little is known about characterization of the *P. adenopoda* genome from chromosome-level genome assembly. In this study, we sequenced and assembled the de novo genome of P. *adenopoda* based on Nanopore sequencing reads, Illumina reads and Hi-C data. This new genome offers insights into tree genome evolution and valuable genomic resources for research in genetic breeding.

## Results and Discussion

### Genome Assembly

Prior to deep sequencing, a genomic survey uncovered the estimated genome size of *P. adenopoda* is ∼423 Mb, and the estimated heterozygosity rate is ∼0.6%. We produced 168× coverage of Nanopore long-read sequencing data, 303× coverage of short reads of Illumina sequencing, and 147× coverage of Hi-C paired-end reads. After primary assembly, comparison, correction, polishing, scaffolding, and removing redundant sequences as well as potential contaminated sequences, a final assembly of 383.43 with contig N50 size of 8.3 Mb was obtained (Table 1). The assembled sequences were further anchored to genetic maps with 19 linkage groups (pseudo-chromosomes) based on Hi-C read pairs. The final assembly consists of 112 scaffolds, which spans 382.29 Mb in total, and with 99.7% anchoring to the linkage groups, including 19 pseudo-chromosomes with sizes ranging from 12.5 to 50.2 Mb (Table 2). The high completeness of this assembly was evidenced by a 97.7% Benchmarking Universal Single Copy Orthologs (BUSCO) recovery score, confirming the continuity and quality of the assembled genome (Table 3).

**Table 1.** Statistics of assembled contigs for *P. adenopoda*.

| Type | Length | Number |
| --- | --- | --- |
| Total | 383,425,513 | 109 |
| Longest | 20,162,781 | - |
| N50 | 8,297,885 | 14 |
| N60 | 5,186,414 | 20 |
| N70 | 3,855,153 | 29 |
| N80 | 2,785,178 | 41 |
| N90 | 1,857,042 | 57 |

**Table 2.**
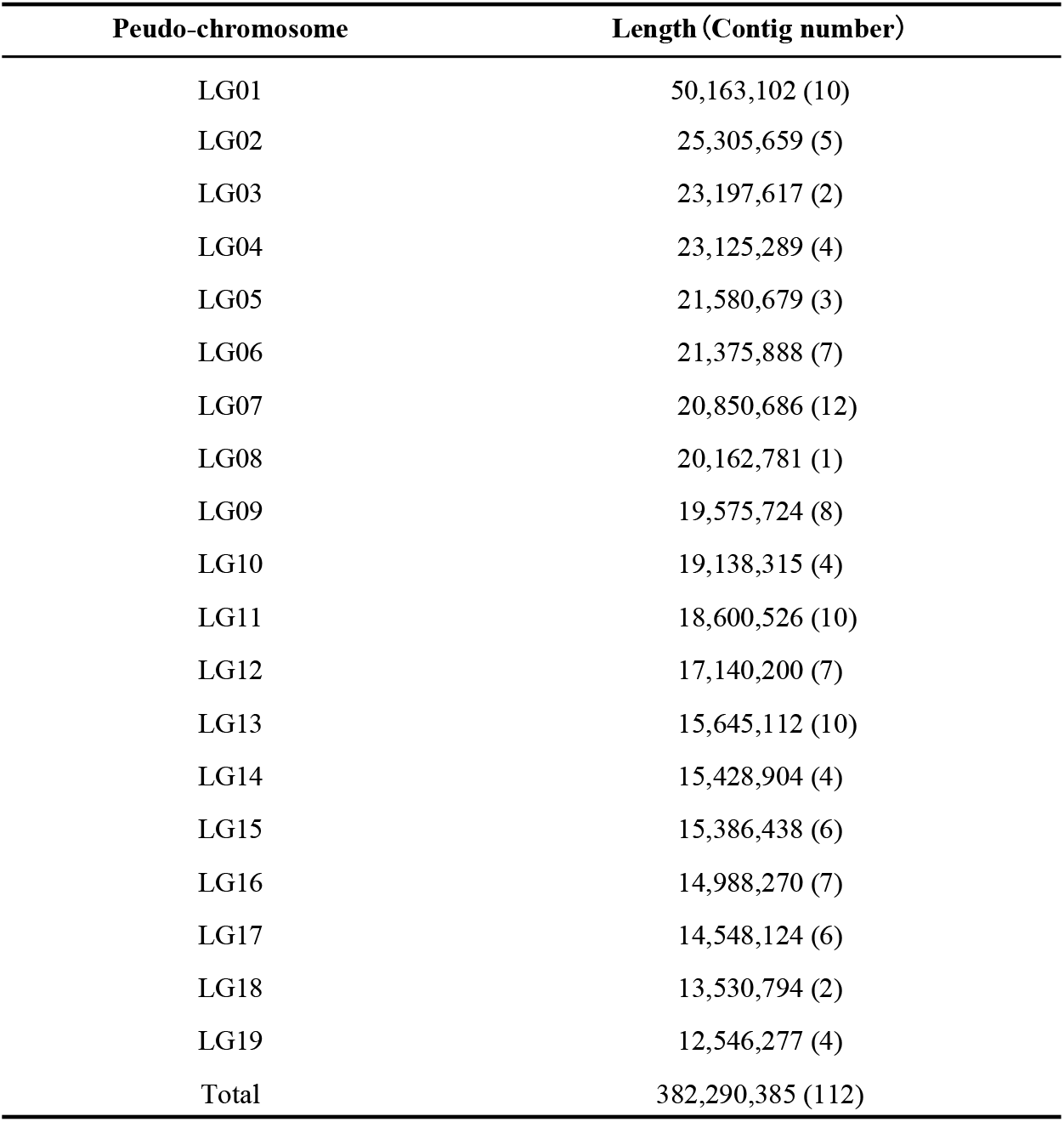
Scaffolding of contigs based on Hi-C data.

**Table 3.** Statistics of BUSCO evaluation for genome assembly.

| BUSCO | Number | Percentage |
| --- | --- | --- |
| Complete BUSCOs (C) | 1,344 | 97.75% |
| Complete and single-copy BUSCOs (S) | 1,115 | 81.09% |
| Complete and duplicated BUSCOs (D) | 229 | 16.65% |
| Fragmented BUSCOs (F) | 8 | 0.58% |
| Missing BUSCOs (M) | 23 | 1.67% |
| Total BUSCO groups searched | 1,375 | 100.00% |

### Genome annotation

Through a combination of de novo and homology-based approaches which integrated in EDTA pipeline, we found that 136.8Mb (35.69%) of the *P. adenopoda* genome is composed of transposable elements (TE), whereas long-terminal repeat (LTR) retrotransposons are the most abundant TEs, representing 17.86% of the *P. adenopoda* genome (Table 4).

**Table 4.** Statistics of repeat sequences in *P. adenopoda* genome.

| TE type | Total size (bp) | Percentage of genome (%) |
| --- | --- | --- |
| **Class I: Retrotransposon** | 69,271,501 | 18.07 |
| **LTR Retrotransposon** | 68,480,792 | 17.86 |
| *Copia* | 13,421,394 | 3.50 |
| *Gypsy* | 38,313,441 | 9.99 |
| unkonwn | 16,745,957 | 4.37 |
| **LINE** | 633,270 | 0.17 |
| **Class II: DNA transposon** | 50,092,626 | 13.07 |
| Helitron | 29,341,144 | 7.65 |
| CACTA | 6,963,805 | 1.82 |
| Mutator | 9,097,395 | 2.37 |
| PIF_Harbinger | 1,094,596 | 0.29 |
| Tcl_Mariner | 468,627 | 0.12 |
| hAT | 3,127,059 | 0.82 |
| **Unclassified** | 17,461,521 | 4.55 |
| **Total** | 136,825,648 | 35.69 |

Combining *ab initio* prediction, homology-based searches, and transcriptome-based approaches, we identified a total of 33,505 protein-coding genes, of which 99.88% (33,465 genes) were located on chromosomes and only 0.12% (40 genes) were annotated on scaffolds. The average gene size is 3,937 bp, with 5.3 exons per gene (Table 5). Functional annotation confirmed that 96.82% of these genes could be assigned to at least one of the databases TrEMBL, Swiss-Prot, NR, Pfam, Interproscan, GO or KEGG (Table 6).

**Table 5.** Statistics of protein-coding genes.

| Feature | number/length |
| --- | --- |
| Gene number | 33,505 |
| Average gene length (bp) | 3936.55 |
| Mean exons number per mRNA | 5.26 |
| Mean CDS number per mRNA | 5.12 |
| Average CDS length (bp) | 1203.06 |
| Average intron length (bp) | 2136.66 |

**Table 6.** Statistics of protein-coding gene functional annotation.

| Database | Number | Percentage (%) |
| --- | --- | --- |
| Pfam | 25,123 | 74.98 |
| Interproscan | 31,328 | 93.53 |
| KEGG | 10,952 | 32.69 |
| NR | 31,647 | 94.45 |
| Swiss-Prot | 25,852 | 77.16 |
| KOG | 29,244 | 87.28 |
| COG | 12,310 | 36.74 |
| Tremble | 32,052 | 95.66 |
| GO | 26,266 | 78.39 |
| Unannotated | 1,067 | 3.18 |

## Materials and Methods

### Plant materials and genome sequencing

Fresh leaf tissues of *P. adenopoda* were collected from wild in Guangyuan, Sichuan, China. Genomic DNA was isolated by CTAB method. For *P. adenopoda* genome, the genomic libraries with 20 Kbp insertions of long-read were constructed and sequenced utilizing the PromethION (ONT) platform of Oxford Nanopore technologies. Paired-end libraries with an average insert length of 350 bp were constructed and sequenced on Illumina HiSeq X Ten platform. Hi-C libraries were prepared from fresh young leaves of the same *P. adenopoda* accession following the standard protocol and sequenced on an Illumina NovaSeq platform. To assist gene annotation, total RNAs from fresh young leaves were extracted with CTAB (Rogers and Bendich 1985) procedure to prepare RNA-seq libraries, which were sequenced on Illumina HiSeq X Ten platform.

### Genome assembly and pseudo-chromosome construction

The Illumina short reads were first used to estimate the genome size of *P. adenopoda* via a 17-bp k-mer frequency analysis with Jellyfish (v2.3.0)^1^. NextDenovo (v2.0-beta.1, https://github.com/Nextomics/NextDenovo) was then used for the preliminary sequence assembly based on the Nanopore long reads. The raw long reads were first error-corrected via NextCorrect with parameters “reads_cutoff=1k, seed_cutoff=30k”, and then assembled via NextGraph with default parameters. To improve the quality of the assembly, corrected ONT long reads (three rounds) and cleaned Illumina short reads (four rounds) were used to polish assembly by Racon v1.3.1^2^ and Nextpolish v1.0.5^3^, separately. The redundant sequences were subsequently removed by using perge_haplotigs v1.1.1^4^ and the obtained genome assemblies were checked for DNA contamination by searching against the NCBI non-redundant nucleotide database (Nt) using BLASTN, with an E-value cutoff of 1e-5. Then, BUSCO (v4.0.5, embryophyta_odb10 download at 16-Oct-2020)^5^ with default settings was applied to the assessment of assembly integrity.

The draft assembly was further scaffolded using Hi-C reads. Briefly, the Hi-C reads were filtered by fastp v0.20.0^6^ with same parameters described above. The clean reads were then aligned into the draft assembled sequences using bowtie2 v2.3.2^7^ with parameters ‘-end-to-end, -very-sensitive -L 30’. The mapped Hi-C reads were processed to obtain the valid reads pairs by HiC-Pro v2.11.4^8^. Scaffolds were anchored into 19 pseudo-chromosomes using LACHESIS^9^ with parameters CLUSTER MIN RE SITES=100, CLUSTER MAX LINK DENSITY=2.5, CLUSTER NONINFORMATIVE RATIO=1.4, ORDER MIN N RES IN TRUNK=60, ORDER MIN N RES IN SHREDS=60, and then followed by manual correction.

### Repeat and gene annotation

Repeat elements were identified by running the pipeline Extensive *de-novo* TE Annotator (EDTA v1.9.3^10^). RepeatMasker v4.1.0^11^ masked the whole genome sequences by using the EDTA-constructed TE library before gene prediction.

An integrated strategy that combined homoeologous gene evidence, transcriptomic supports and *ab initio* predictions was applied to predict protein-coding genes in the *P. adenopoda* genome. For homology-based gene prediction, Genewise v2.4.1^12^ was used to predict gene models based on published protein sequences of six species (*Populus trichocarpa, Populus euphratica, Salix brachista, Salix purpurea, Arabidopsis thaliana* and *Vitis vinifera*). For transcriptome-based prediction, the clean RNA-seq reads were firstly aligned to the *P. adenopoda* genome using HISAT v2.2.1^13^, and then assembled to transcripts using Trinity v2.8.4^14^ with default parameters. These assembled transcripts were subsequently aligned to the corresponding genome to predict gene structure using PASA v2.4.1^15^. For ab initio gene prediction, AUGUSTUS v3.3.2^16^ was employed with default parameters after incorporating the transcriptome-based and homology-based evidence for gene model training. All candidate genes models generated by three approaches were combined and integrated by EvidenceModeler v1.1.1^17^ to obtain a non-redundant consensus gene set. Untranslated regions and alternative transcripts were predicted by PASA v2.4.1 to further correct the predicted genes.

Functional annotation for predicted genes was achieved by using a blast homologue search against public protein databases, including NCBI nonredundant protein database (NR), Swiss-Prot, TrEMBL, COG and KO. Functional motifs and domains of each gene were identified by InterProScan (v5.32-71.0)^18^. Gene ontology (GO) terms and KEGG Automatic Annotation Server was utilized to explore the KEGG pathways for each gene were assigned by InterProScan and KEGG Automatic Annotation Server, respectively.

